# The FLI portion of EWS/FLI contributes a transcriptional regulatory function that is distinct and separable from its DNA-binding function in Ewing sarcoma

**DOI:** 10.1101/2020.10.29.355859

**Authors:** Megann A. Boone, Cenny Taslim, Jesse C. Crow, Julia Selich-Anderson, Iftekhar A. Showpnil, Benjamin D. Sunkel, Meng Wang, Benjamin Z. Stanton, Emily R. Theisen, Stephen L. Lessnick

## Abstract

**Background:** Ewing sarcoma is an aggressive bone cancer in children and young adults that contains a pathognomonic chromosomal translocation: t(11;22)(q24;q12). The encoded protein, EWS/FLI, fuses the low-complexity amino-terminal portion of EWS to the carboxyl-terminus of FLI. The FLI portion contains an ETS DNA-binding domain and adjacent amino- and carboxyl-regions. Early studies using non-Ewing sarcoma cellular models provided conflicting information on the role of these adjacent regions in the oncogenic function of EWS/FLI. We therefore sought to define the specific contributions of each FLI region to EWS/FLI activity in an appropriate Ewing model, and in doing so, to better understand Ewing sarcoma development mediated by the fusion protein.

**Methods:** We used a “knock-down/rescue” system to replace endogenous EWS/FLI expression with mutant forms of the protein in Ewing sarcoma cells and tested these for oncogenic transformation using soft-agar colony forming assays. These data were complemented by DNA-binding assays using fluorescence anisotropy, genomic localization assays using CUT&RUN, transcriptional regulation studies using luciferase reporter assays and RNA-sequencing, as well as chromatin accessibility assays using ATAC-sequencing.

**Results:** We found that the DNA-binding domain and short flanking regions of FLI were required for oncogenic transformation, gene expression, genomic localization and chromatin accessibility when fused to the amino-terminal EWS-portion from EWS/FLI, but that the remaining regions of FLI were dispensable for these functions. Removal of a carboxyl-terminal alpha-helix from the short flanking regions of the DNA-binding domain of FLI created a hypomorphic EWS/FLI that retained normal DNA binding, genomic localization, and chromatin accessibility, but had significantly restricted transcriptional activity and a near total loss of oncogenic transformational capacity.

**Conclusions:** The DNA-binding domain and carboxyl-terminal short flanking region of FLI are the only portions of FLI required for EWS/FLI-mediated oncogenic transformation in a Ewing sarcoma cellular context. In addition to the well-defined DNA-binding function of FLI, this additional alpha-helix immediately downstream of the DNA-binding domain contributes a previously-undescribed function in gene regulation and oncogenic transformation. Understanding the function of this critical region could provide new therapeutic opportunities to target EWS/FLI in Ewing sarcoma.

## Background

Ewing sarcoma is a bone-tumor of children and young adults (2). These tumors always contain chromosomal translocations that encode fusions between a member of the FET protein family and one of the ETS transcription factors (1, 3). The most common translocation (in ~85% of patients) is the t(11;22)(q24;q12), which fuses *EWSR1* to *FLI1* (1, 4, 5). The *EWSR1/FLI1* fusion encodes the EWS/FLI protein (3, 6). Multiple studies have demonstrated that EWS/FLI has oncogenic function and serves as the driver oncoprotein in Ewing sarcoma (1, 4, 7). Indeed, EWS/FLI is often the only genetic abnormality in these otherwise “genomically-quiet” tumors (8). Thus, determining the mechanisms underlying the oncogenic function of EWS/FLI is critical to understanding Ewing sarcoma tumorigenesis, identifying new therapeutic approaches for this aggressive disease, and may also shed light on the oncogenic mechanisms of other “ETS-associated” tumors.

EWS/FLI functions as an aberrant transcription factor and dysregulates several thousand genes (9, 10). The intrinsically-disordered low-complexity domain of EWS contributes strong transcriptional activating and repressing functions to the fusion (11–13). The mechanisms by which the EWS-portion mediates these functions are only beginning to be understood, but may include the recruitment of epigenetic co-regulators and RNA-polymerase II, perhaps via the formation of transcriptional “hubs” consisting of low-affinity/high-valency interactions, phase-separated droplets, or even polymerized fibrils (9, 14–17).

FLI is a member of the ETS family of transcription factors (18–20). The 28 ETS family members in humans are defined by the presence of highly conserved winged helix-turn-helix DNA-binding domains (DBD) (18). The preferred high-affinity (HA) binding sequence for FLI is “ACCGGAAGTG”, while other family members bind similar sequences containing a GGA(A/T) core surrounded by additional base pairs (18, 21). Structural studies have demonstrated that the third alpha-helix of the FLI DNA-binding domain (part of the winged helix-turn-helix structure) binds in the major groove of DNA (19, 22, 23). Several ETS family members demonstrate “autoinhibitory” activity in which domains adjacent to the DNA-binding domain reduce the ability of ETS factors to bind DNA (24). However, FLI harbors only minimal autoinhibitory activity (~2-3 fold reduction in binding affinity) (24, 25). In addition to binding classic ETS HA sites, EWS/FLI also binds to microsatellite sequences consisting of multiple “GGAA” tetrameric repeats (26–28). There are thousands of GGAA-microsatellite sequences scattered throughout the human genome, and many of these serve as EWS/FLI-response elements associated with target genes critical for oncogenic transformation in Ewing sarcoma (26–28). The ability of EWS/FLI to regulate target genes through GGAA-microsatellites appears to be a neomorphic function gained by the fusion protein as compared to wild-type FLI. Along with the ETS DNA-binding domain, the FLI portion of the fusion contains additional amino-terminal and carboxyl-terminal regions of uncertain function.

The cell of origin of Ewing sarcoma is not known (29). Early studies that analyzed the role of the FLI portion of EWS/FLI used heterologous cell types, such as NIH3T3 murine fibroblasts, with conflicting results (29). For example, expression of wild-type EWS/FLI induces oncogenic transformation of NIH3T3 cells, while expression of an EWS/FLI mutant harboring a complete deletion of the ETS DNA-binding domain did not, demonstrating a critical role for DNA-binding in the function of EWS/FLI (7). In contrast, a later study demonstrated that a partial deletion of the ETS DNA-binding domain (that also disrupted DNA-binding) instead retained the capability of inducing oncogenic transformation of NIH3T3 cells (30). Subsequent studies in patient-derived Ewing sarcoma cells showed that a DNA-binding defective mutant of EWS/FLI was unable to mediate oncogenic transformation in cells, thus proving that DNA-binding is absolutely required for EWS/FLI-mediated transformation in a more relevant Ewing cellular model (13). The carboxyl-terminal region of FLI (outside of the DNA-binding domain) was also evaluated in the NIH3T3 model and determined to be important for transcriptional control and oncogenic transformation mediated by EWS/FLI, though this has not been reproduced in a Ewing sarcoma model (31). Later work demonstrated that gene expression patterns mediated by EWS/FLI in the NIH3T3 model were drastically different from those in Ewing sarcoma cellular models, suggesting that EWS/FLI may utilize alternative mechanisms to drive oncogenesis in different systems and that model system selection is important (29). To our knowledge, a systematic evaluation of the FLI portion of EWS/FLI in Ewing sarcoma cells has yet to be reported and so the roles of various regions of FLI in EWS/FLI-mediated oncogenic transformation remain unknown.

To address this important gap in knowledge, we now report an analysis of the FLI portion of EWS/FLI in Ewing sarcoma cells using our well-validated “knock-down/rescue” system. This model allowed us to identify an uncharacterized region just outside of the DNA-binding domain of FLI that is absolutely essential for EWS/FLI-mediated transcriptional regulation and oncogenic transformation. Furthermore, this system allowed us to explore the mechanistic contributions of this region using luciferase reporter and whole genome RNA-sequencing assays, DNA-binding assays and genomic localization studies, and evaluation of open chromatin regions using ATAC-sequencing. These studies demonstrate a unique contribution of this region in mediating gene expression and subsequent oncogenic transformation that is independent of DNA-binding or the modulation of open chromatin states.

## Methods

### Constructs and retroviruses

Puromycin-resistant retroviral vectors encoding shRNAs targeting *Luciferase* (iLuc) or the 3’-UTR of endogenous EWS/FLI mRNA (iEF) were previously described (28, 32). Full-length EWS/FLI and mutants (all containing amino-terminal 3xFLAG-tags) were cloned into pMSCV-Hygro (Invitrogen) with sequence details provided in Additional file 1. Luciferase reporter constructs (in pGL3 vectors; Promega Corporation) were previously described (28). The FLI DBD or FLI DBD+ proteins (each containing a carboxyl-terminal 6xHistidine tag) were expressed using pET28a plasmids (EMD Chemicals).

### Cell culture methods

HEK-293EBNA (Invitrogen) and A673 cells (ATCC) were grown, retroviruses produced and used for infection, and soft agar assays were performed as described (28, 32, 33). STR profiling and mycoplasma testing are performed annually on all cell lines. Dual luciferase reporter assays were performed in HEK-293EBNA cells as previously described (28).

### Immunodetection

Whole-cell or nuclear protein extraction, protein quantification, and Western blot analysis was performed as previously described (28, 32, 33). Immunoblotting was performed using anti-FLAG M2 mouse (Sigma F1804-200UG), anti-α-Tubulin (Abcam ab7291), and anti-Lamin B1 (Abcam ab133741). Membranes were imaged using the LiCor Odyssey CLx Infrared Imaging System.

### qRT-PCR

Total RNA was extracted from cells using the RNeasy Extraction Kit (Qiagen 74136). Reverse transcription and qPCR were performed using the iTaq Universal SYBR Green 1-Step Reaction Mix (BioRad 1725151) on a Bio-Rad CFX Connect Real-Time System. Primer sequences are found in Additional file 2.

### Recombinant protein purification

Recombinant 6xHistidine-tagged FLI DBD and FLI DBD+ protein was purified as previously described (28).

### Fluorescence anisotropy

Fluorescence anisotropy was performed using recombinant protein and fluorescein-labeled DNA duplexes as previously described using sequences provided in Additional file 3 (28).

### CUT&RUN and Analysis

Two biological replicates for each knock-down/rescue sample were analyzed by CUT&RUN using the anti-FLAG M2 mouse antibody (Sigma F1804-200UG) as described and sequenced with the Illumina HiSeq4000 (32). Raw reads were trimmed, de-duplicated, aligned to hg19 reference genomes, and peaks were called using macs2 and DiffBind (Bioconductor) using I”EF + Empty Vector” samples as controls (34). Bigwig files combining two replicates with normalization option “RPGC” were created using Deeptools (35). Overlapping peak analysis was completed using R packages ChIPpeakAnno and Genomic Ranges (36, 37).

### RNA-sequencing and Analysis

RNA-sequencing was performed on two biological replicates each for knock-down/rescue A673 sample. TruSeq Stranded mRNA Kit (Illumina Cat. No. 20020594) was used to prepare cDNA libraries from total RNA and sequenced on Illumina HiSeq4000 to generate 150-bp paired-end reads. Reads were analyzed for quality control, trimmed, aligned to the human genome and analyzed for differential analysis (using FASTQC, Multiqc, Trim_galore, STAR version 2.5.2b, DESeq2) (38). GSEA (Version 4.0.3) analysis was performed on RNA-sequencing data (39). Significantly activated and repressed genes were defined using a log2(FC) of |1.5| cutoff for EF and EF DBD+ to create gene sets. EF DBD genes were used as the rank-ordered gene list to compare with these gene sets. RNA-expression scatterplot analysis was performed as previously described (32).

### ATAC-sequencing and Analysis

ATAC-sequencing was performed on two separate biological replicates for knock-down/rescue A673 cells as previously described and sequenced with Illumina HiSeq4000 (40, 41). The ENCODE pipeline was used for trimming, alignment to hg19 reference genome, and peak calling on individual replicates (ENCODE Project). RegioneR was used to perform permutation test and test significance of overlapping ATAC peaks in different samples (37). EnrichedHeatmap, ggplot2, ChIPpeakAnno, and GenomicRanges were used to calculate overlapping regions and create heatmaps (36, 37, 42, 43). Differential ATAC peak analysis was completed using DiffBind (Bioconductor) and DESeq2 with an FDR<0.05 (38).

### Statistical Analysis

Luciferase assay, soft agar assay, and PCR data are presented as mean ± SEM. Fluorescence anisotropy data are presented as mean ± standard deviation. Significance of experimental results was determined using a Student’s t-test for comparison between groups. P-values less than 0.05 were considered to be significant.

## Results

### Amino- and carboxyl-terminal regions of FLI are dispensable for EWS/FLI-mediated transcriptional activation in luciferase reporter assays

We first sought to determine the role of the amino- and carboxyl-regions of FLI in EWS/FLI-mediated transcriptional activation using a well-defined luciferase reporter assay containing a 20xGGAA-microsatellite response element and a minimal promoter (28). We used the “type IV” EWS/FLI fusion (containing regions encoded by exons 1-7 of *EWSR1* fused to exons 7-9 of *FLI1*) as the full-length protein (containing the 3xFLAG-tag), as this has been extensively used in our previous studies (28, 32). We created 3xFLAG-tagged constructs to express “EF ΔN-FLI” and “EF ΔC-FLI” mutants that harbor deletions in the FLI portion of the fusion that were either amino-terminal to or carboxyl-terminal to the FLI DNA-binding domain, respectively (Figure 1A). EF ΔC-FLI is similar to the Δ89-C protein that showed diminished oncogenic potential in the NIH3T3 model system (31). Expression plasmids encoding these proteins were co-transfected with the 20xGGAA-microsatellite luciferase reporter into HEK-293EBNA cells. All three proteins were expressed at similar levels (Figure 1B). We found that all three versions of EWS/FLI were capable of activating luciferase reporter gene expression to similar levels (Figure 1C). These data demonstrate that neither the amino-terminal nor the carboxyl-terminal region of FLI is required for transcriptional activation mediated by EWS/FLI in this system.

**Figure 1.**
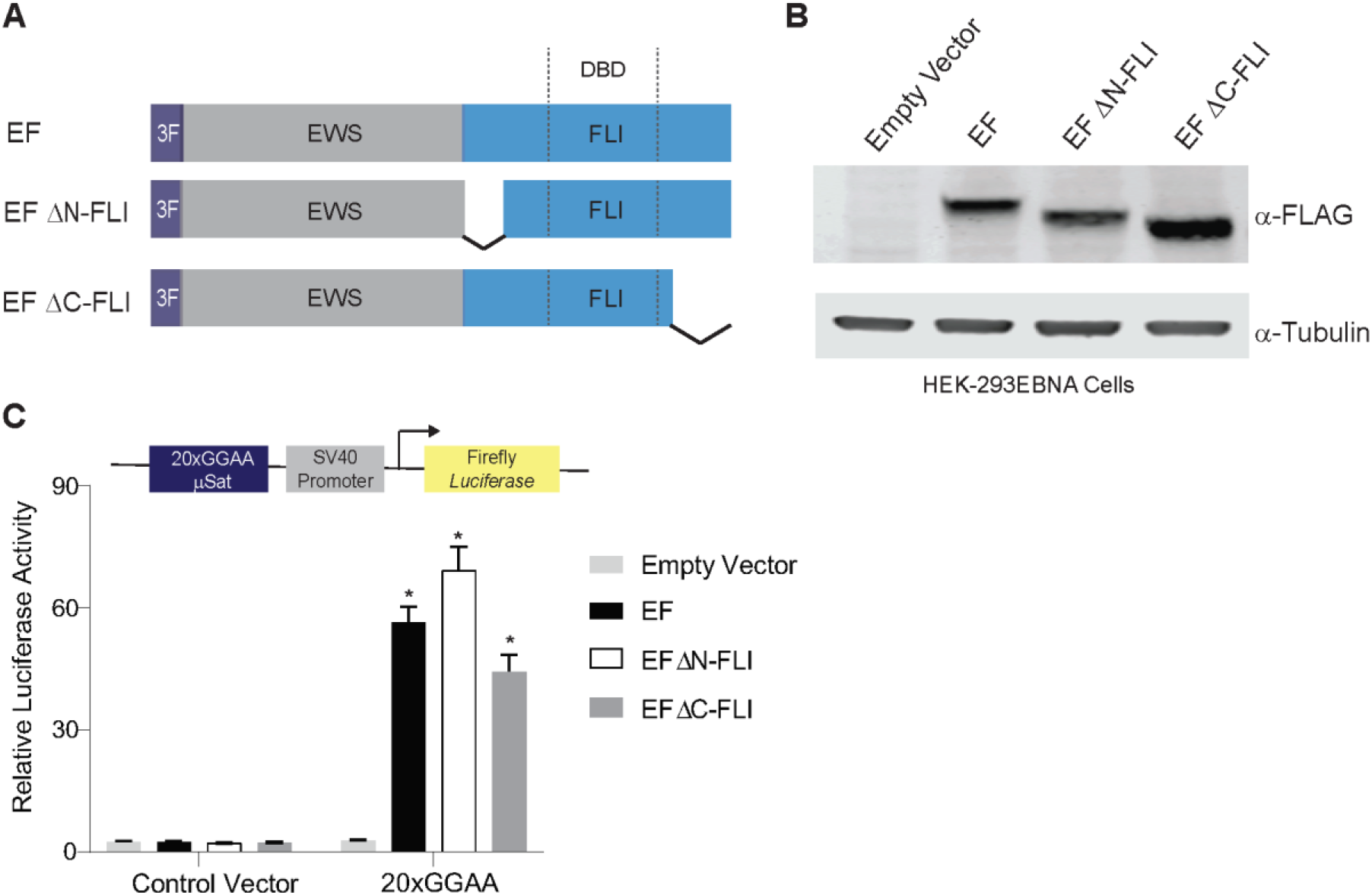
Amino- and carboxyl-terminal regions of FLI are not required for EWS/FLI-mediated transcriptional activation. (A) Protein schematic of 3xFLAG-tagged (3F) EWS/FLI (EF) cDNA constructs with FLI deletions. EWS is represented in grey, FLI is represented in blue, and dashed lines in the FLI portion represent the 85-amino acid ETS DNA-binding domain (DBD) of FLI. In each construct, EWS is fused directly to the FLI portion, but connecting lines are shown here to represent regions of FLI that are eliminated in each construct. EF represents a full-length “type IV” EWS/FLI translocation. EF ΔN-FLI and EF ΔC-FLI indicate constructs where EWS was fused to a version of FLI with a deletion in the N- or C-terminal region of, respectively. (B) Western blot of 3xFLAG-tagged EWS/FLI protein expression in HEK-293EBNA cells. Membranes were probed with either α-FLAG or α-tubulin antibodies. (C) Dual luciferase reporter assay results for the indicated cDNA constructs co-transfected into HEK-293EBNA cells with a Control Vector harboring no GGAA-repeats, or a vector containing 20xGGAA-repeats (shown schematically above the graph). Data are presented as mean ± SEM (n=6 independent replicates). Asterisks indicate that the activity of EF, EF ΔN-FLI, and EF ΔC-FLI are each statistically significant when compared to Empty Vector on a 20xGGAA μSat (p-value <0.05). The activity of EF ΔN-FLI and EF ΔC-FLI are not statistically different from EF on the 20xGGAA μSat (p-value = 0.8).

### Regions immediately adjacent to the DNA-binding domain of FLI are required for oncogenic function of EWS/FLI in a Ewing sarcoma cellular model

We next hypothesized that the only critical region in the FLI portion of EWS/FLI is the ETS DNA-binding domain itself. The ETS DNA-binding domain of FLI is not well-defined in the published literature. The ETS domain is often referred to as an 85-amino acid sequence (18, 19, 21). However, other structural and functional studies of FLI defined a larger region of FLI as the ETS domain that included short amino- and carboxyl-extensions to the 85-amino acid “core” (7, 44). To test both possible “ETS domains”, we created two new mutant forms of EWS/FLI: “EF DBD” that fuses EWS directly to the 85-amino acid ETS domain and “EF DBD+” that fused EWS to a 102-amino acid ETS DNA-binding domain (containing 7- and 10-amino acid extensions on the amino-terminal and carboxyl-terminal sides of DBD, respectively) that our laboratory has used in prior studies (Figure 2A) (26).

**Figure 2.**
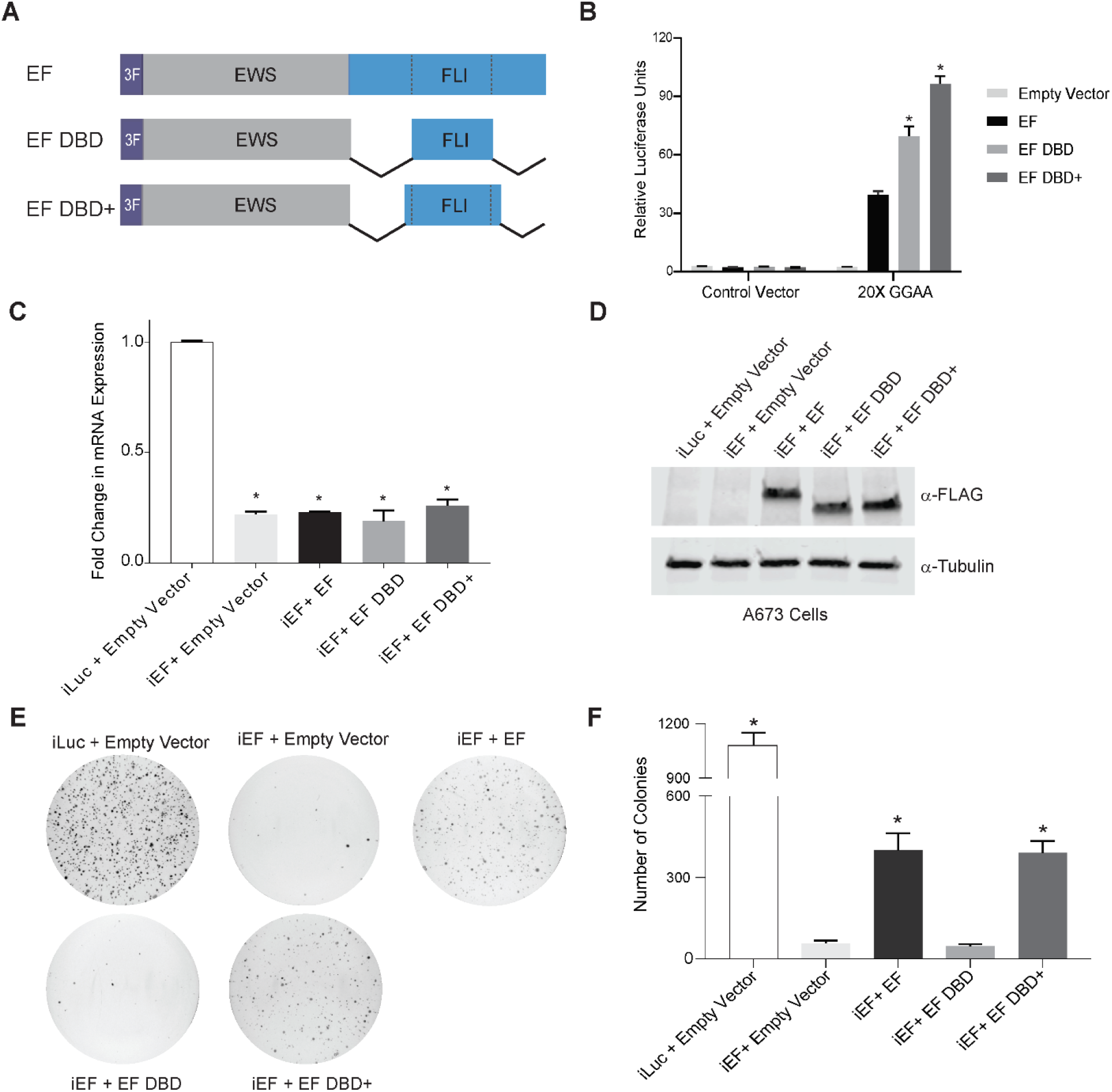
Oncogenic transformation capacity of EWS/FLI affected by short regions surrounding the FLI DBD. (A) Protein schematic of 3xFLAG-tagged (3F) EWS/FLI cDNA constructs with deleted FLI domain regions. EF represents a full-length type IV EWS/FLI, EF DBD represents EWS fused directly to the 85-amino acid DNA-binding domain of FLI, and EF DBD+ represents EWS fused to a 102 amino acid region of FLI that contains the 85 amino-acid DNA-binding domain with 7 additional amino-acids on the amino-terminal side and 10 additional amino-acids on the carboxyl-terminal side (B) Dual luciferase reporter assay results for the indicated constructs tested on control and 20xGGAA μSat-containing plasmids (as described in Figure 1). Data are presented as mean ± SEM (n=6 independent replicates). Asterisks indicate that the activity of EF DBD and EF DBD+ are each statistically higher than EF (p-value < 0.001). (C) Representative qRT-PCR results of endogenous EWS/FLI in A673 cells harboring the indicated constructs (iLuc is a control shRNA targeting luciferase and iEF is an shRNA targeting the 3’UTR of endogenous EWS/FLI; n=1). EWS/FLI mRNA value are normalized to RPL30 mRNA control values. Asterisks indicate samples are statistically different as compared to control iLuc + Empty Vector cells (p-value < 0.001). (D) Western blot analysis of exogenous EWS/FLI protein expression corresponding to samples shown in panel C. Protein constructs were detected using α-FLAG antibody and α-Tubulin was used as a loading control. (E) Representative soft agar assay results of A673 Ewing sarcoma cells containing the indicated constructs. (F) Soft agar assay colony formation quantification. Data presented as mean ± SEM (n=9 independent replicates). Asterisks indicate p-value <0.001 as compared to iEF + Empty Vector.

Similar levels of protein expression were observed following transfection into HEK-293EBNA cells (Supplemental Figure 1A). Luciferase reporter assays using the 20xGGAA-microsatellite response element revealed that both EF DBD and EF DBD+ induced robust transcriptional activation and were even more active than full-length EWS/FLI itself (EF) (Figure 2B, p-value<0.001).

To determine whether the results observed using the luciferase reporter assay would translate to a more relevant Ewing sarcoma cellular model, we used our “knock-down/rescue” system to replace endogenous EWS/FLI with exogenous constructs in patient-derived A673 Ewing sarcoma cells (45). Briefly, retrovirally-expressed shRNAs targeting a control gene (iLuc) or the 3’-UTR of endogenous EWS/FLI (iEF) were introduced and efficient reduction of endogenous EWS/FLI mRNA was achieved (Figure 2C). Exogenous expression of EWS/FLI was “rescued” through retroviral expression of cDNA constructs (EF, EF DBD, and EF DBD+) lacking the endogenous 3’-UTR (Figure 2D).

These “knock-down/rescue” cells were seeded into soft agar to measure anchorage-independent colony formation as a measure of oncogenic transformation (Figure 2E-F). Positive control cells (iLuc + Empty Vector) showed high rates of colony formation, while cells with diminished levels of EWS/FLI (iEF + Empty Vector) showed a near total loss of transformation capacity that was rescued by re-expression of full-length EWS/FLI (iEF + EF). (Figure 2E-F). Interestingly, expression of EF DBD+ (iEF + EF DBD+) rescued colony formation to the same level as full-length exogenous EF, but the smaller EF DBD construct completely failed to rescue colony formation (Figure 2E-F, p-value<0.005). These data define a significant functional difference between EF DBD and EF DBD+ in the A673 Ewing sarcoma model that is not correlated to their transcriptional activity in the luciferase reporter assay.

### DNA-binding and genomic localization of EWS/FLI are nearly identical in FLI domain mutants

The inability of EF DBD to rescue A673 cell colony growth suggested a loss of a critical function as compared to EF DBD+. The only difference between the EF DBD and EF DBD+ constructs is in the 17-amino acids flanking the 85-amino acid DNA-binding domain core. We therefore reasoned that these flanking amino acids may contribute to EWS/FLI DNA-binding affinity. To test this hypothesis, we performed *in vitro* fluorescence anisotropy studies to compare the ability of FLI DBD or FLI DBD+ recombinant protein to bind fluorescein-labeled DNA duplexes (Figure 3A, Supplemental Figure 2A-B). We tested an ETS high affinity (HA) site, a 2xGGAA-repeat microsatellite, and a 20xGGAA-repeat microsatellite (Figure 3B-D). We found that both FLI DBD and FLI DBD+ bound each DNA element with similar dissociation constant (K_D_) values (Figure 3B-D). These data discount the idea that a significant difference in DNA-binding affinity is the underlying defect in EF DBD.

**Figure 3.**
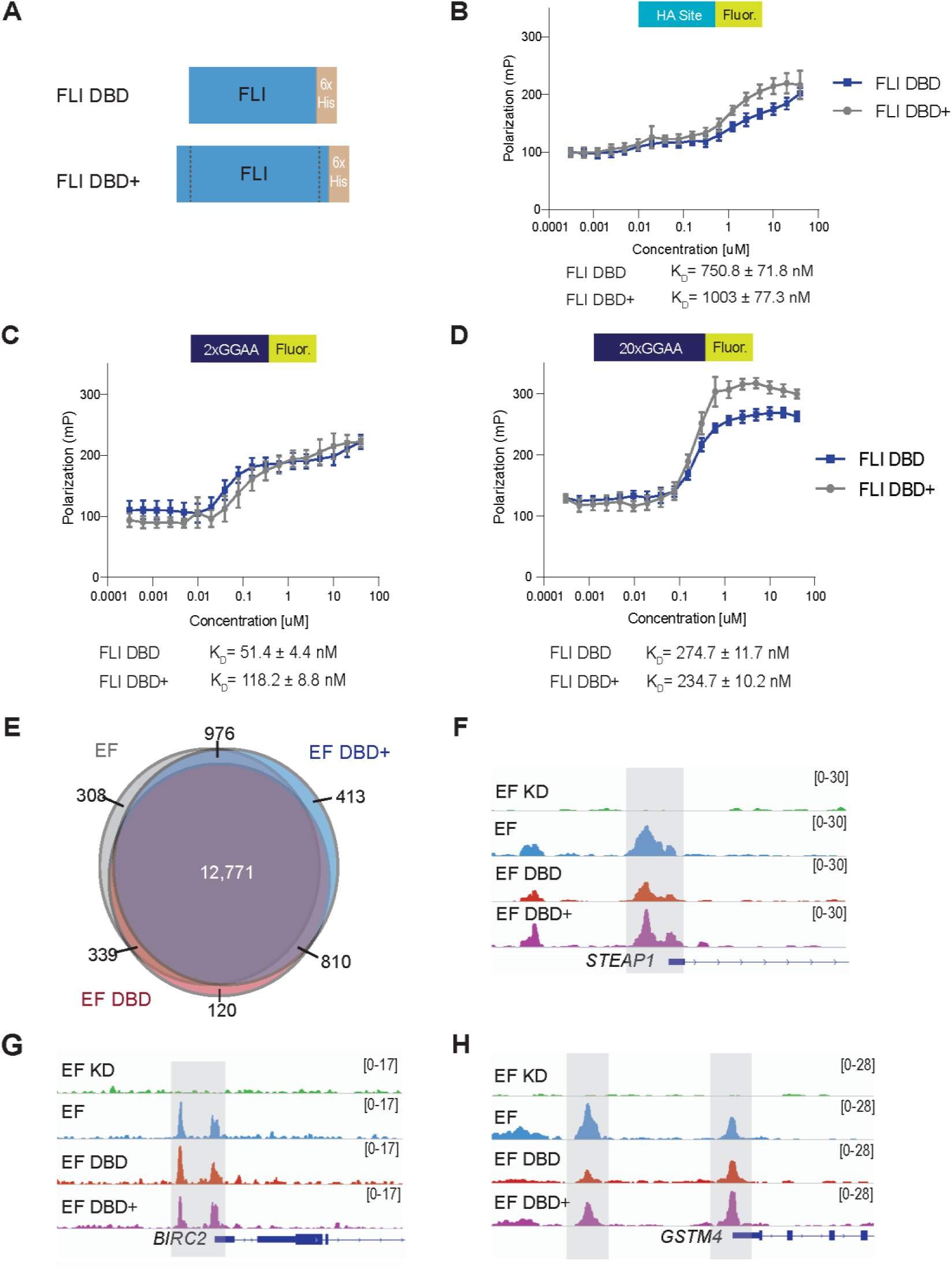
DNA-binding and genomic localization properties of EWS/FLI unaltered by deletions flanking the FLI DNA-binding domain. (A) Protein schematic of FLI DBD or FLI DBD+ recombinant protein (with C-terminal 6xHistidine-tag [6xHis]). (B-D) Fluorescence anisotropy assay results for FLI DBD and FLI DBD+ recombinant proteins (0-40 μM) on the following fluorescein-labeled DNA sequences: (B) ETS high affinity (HA) site DNA, (C) 2x-repeat GGAA μSat DNA, and (D) 20x-repeat GGAA μSat DNA (n=9 independent replicates). Dissociation constants (K_D_) for FLI DBD and FLI DBD+ are noted for each DNA response element. (E) Venn diagram comparing peaks called in CUT&RUN for EWS/FLI construct localization in knock-down/rescue cells (EF = iEF + EF; EF DBD = iEF + EF DBD; EF DBD+ = iEF + EF DBD+) when compared to cells that did not contain a rescue construct (iEF + Empty Vector). The number of peaks overlapping between constructs are indicated on the Venn diagram. (F-H) Representative CUT&RUN peak tracks from IGV are shown for iEF + Empty Vector (EF KD), EF, EF DBD, and EF DBD+ samples. Examples of peaks from EWS/FLI-associated HA-site regulated genes ([F] *STEAP1* and [G] *BIRC2*) and GGAA-μSat-regulated genes ([H] *GSTM4*) are highlighted. Peak track scales are shown on the right.

Although *in vitro* DNA-binding was nearly identical between FLI DBD and FLI DBD+ recombinant proteins, we next considered if differences in DNA-binding may only be revealed in the context of a chromatinized human genome. To assess this possibility, we performed CUT&RUN (Cleavage Under Targets & Release Under Nuclease) to determine the genomic localization patterns of our 3xFLAG-tagged EF, EF DBD, and EF DBD+ proteins in A673 cells using our knock-down/rescue system (32, 46). An anti-FLAG antibody was used to ensure that we evaluated the localization of the exogenous “rescue” constructs and not any low-level residual EWS/FLI remaining after the knock-down. We found that CUT&RUN identified a similar number of binding peaks between full-length EF (14,040), EF DBD+ (14,970), and EF DBD (14,394). Comparison of the binding locations for each construct demonstrated that 90% of EF DBD peaks overlap with those of EF and EF DBD+ (Figure 3E, adjusted p-value < 0.001). Further exploration of EWS/FLI-bound high-affinity sites and microsatellites did not identify any significant differences between EF DBD and EF or EF DBD+ (representative peak tracks are shown in Figure 3F-H). Taken together, these data indicate that there are no large-scale changes in DNA-binding capabilities that might explain the inability of EF DBD to rescue oncogenic transformation in Ewing sarcoma cells.

### EF DBD exhibits a hypomorphic gene regulatory capability in Ewing sarcoma cells

The above studies demonstrated that genome-wide localization is nearly-identical between the full-length and mutant EWS/FLI constructs. Although luciferase assays (Figure 2B) showed strong transcriptional activation by EF DBD, we next considered the possibility that transcriptional regulatory function of EF DBD might be disrupted in a more relevant Ewing sarcoma model. To test this hypothesis, we performed RNA-sequencing on knock-down/rescue cells expressing full-length EWS/FLI (EF), EF DBD+, or EF DBD.

Full-length EWS/FLI (EF) regulated 4,124 genes and EF DBD+ regulated 3,374 genes (at adjusted p-values < 0.05). Importantly, 90% of the genes regulated by EF DBD+ were also regulated by EF. In contrast, EF DBD regulated only 964 genes (adjusted p-value < 0.05). To determine if activated or repressed genes were disproportionately affected, we analyzed each group separately. There was a similar loss of regulation in each group, although the genes that were regulated by EF DBD mostly overlapped with the fully-functional constructs (Figure 4A-B).

**Figure 4.**
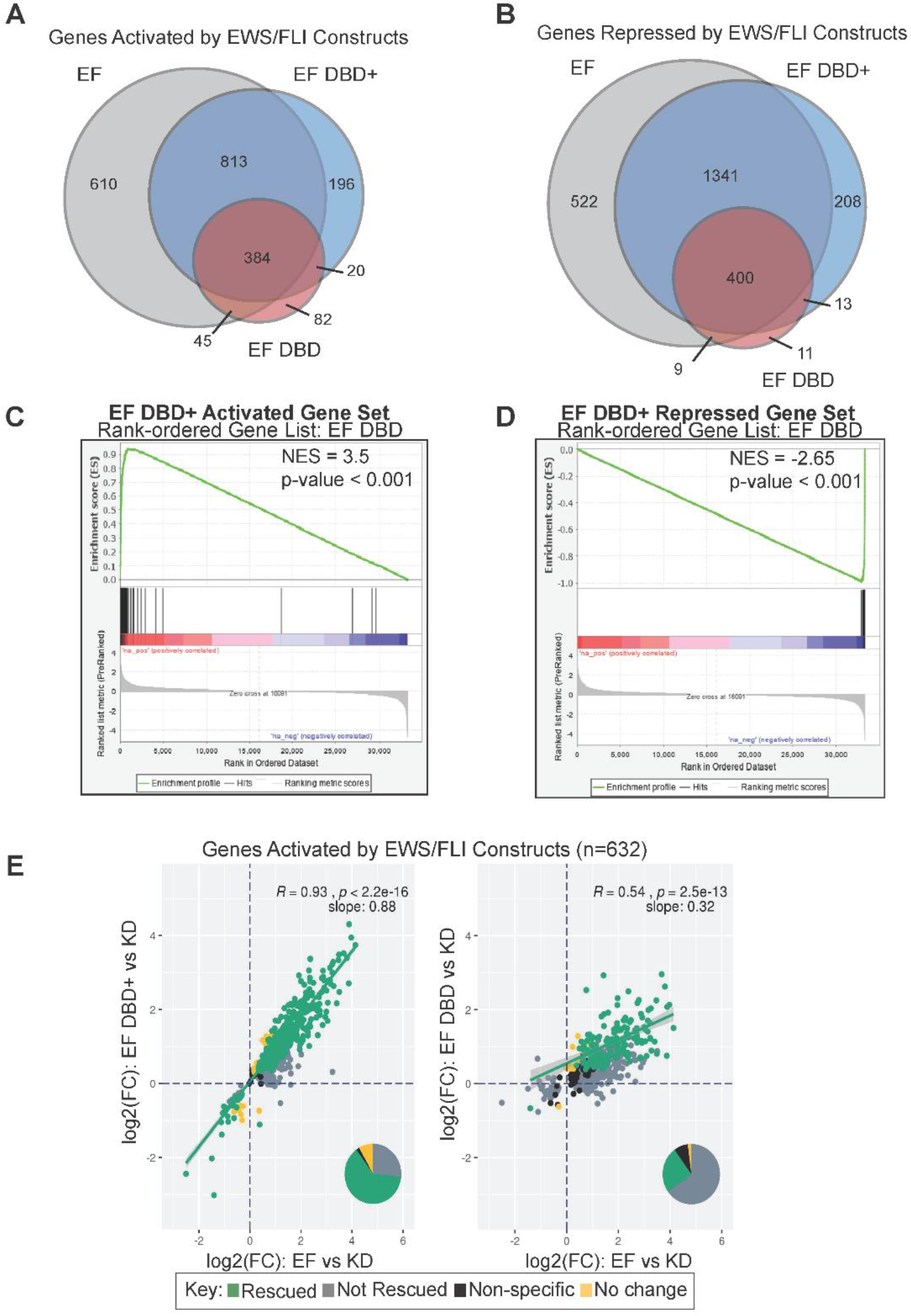
EWS/FLI-driven transcriptional regulation diminished by FLI DBD flanking deletions in Ewing sarcoma cells. (A-B) Venn diagram analysis of RNA-sequencing data comparing genes significantly (A) activated or (B) repressed in A673 cells rescued with the indicated constructs (full-length EWS/FLI [EF], EF DBD, and EF DBD+) when compared to A673 cells with no exogenous EWS/FLI construct (iEF + Empty Vector) (adjusted p-value (FDR) < 0.05). (C-D) GSEA analysis comparing all genes regulated by EF DBD as the rank-ordered gene list to (C) genes activated by EF DBD+ (log2(FC) > 1.5, FDR < 0.05) as the gene set or (D) genes repressed by EF DBD+ (log2(FC) < −1.5, FDR < 0.05) as the gene set. (E) Genes significantly activated by endogenous EWS/FLI were defined using a previous RNA-sequencing dataset (47). Genes activated by EF, EF DBD, and EF DBD+ were compared to this list of EWS/FLI-activated genes. Scatterplots comparing genes activated by EF (on the x-axis) to EF DBD+ (left) or EF DBD (right) (on the y-axis) were plotted to determine the ability of these constructs to rescue expression these genes. Significance was defined by a log2(FC) > 0 and an adjusted p-value < 0.05. Pearson correlation coefficient and associated p-values with slope are noted on the plots. Pie charts represent the proportion of genes found in each of the described groups is also depicted.

We next performed a more detailed evaluation of the RNA-sequencing data using Gene Set Enrichment Analysis (GSEA). We asked where the activated and repressed gene sets of EF DBD fall in comparison to the rank-ordered gene expression list of EF DBD+. We found very strong correlations of both the activated and repressed gene sets (|NES| of 3.5 and 2.65, respectively, p < 0.001; Figure 4C-D). Even stronger correlations were observed when EF DBD-regulated gene sets were compared with EF activated and repressed genes (|NES| of 7.09 and 5.65, respectively, p < 0.001; Supplemental Figure 3A-B).

Closer inspection of the GSEA results revealed a near-complete “stacking” of the EF DBD-regulated genes at the furthest edges of the EF DBD+ (or full-length EF) rank-ordered lists. This suggested that EF DBD may only be able to significantly rescue a portion of the EF DBD+ or EF-regulated genes, while other genes are still regulated, though to a lesser extent and are not called as significant. We therefore hypothesized that EF DBD functions as an attenuated, hypomorphic version of EWS/FLI. To test this hypothesis, we performed a scatterplot analysis to compare the ability of these constructs to rescue previously-reported EWS/FLI-regulated genes (47). On the x-axis, we plotted the expression levels of genes regulated by the full-length EF construct (activated genes in Figure 4E and repressed genes in Supplemental Figure 3C). On the y-axis, we plotted the expression levels of genes regulated by EF DBD+ or EF DBD. Transcriptional regulation by EF DBD+ at activated and repressed genes was highly correlated with regulation by EF (slope=0.88 with R=0.93 and slope=0.94 with R=0.97, respectively; p-value < 2.2e-16; Figure 4E left panel, Supplemental Figure 3C left panel). In contrast, EF DBD demonstrated much weaker correlations (slope=0.32 with R=0.54 for activated genes, p-value < 2.5e-13; slope=0.54 with R=0.78 for repressed genes, p-value < 2.2e-16; Figure 4E right panel, Supplemental Figure 3C right panel). As regulation of EF DBD was still correlated with EF, these data suggested that it is regulating a similar set of genes, albeit more weakly than EF or EF DBD+.

Taken together, these data indicate that EF DBD is significantly attenuated in its ability to both up- and down-regulate gene expression in patient-derived Ewing sarcoma cells. Thus, EF DBD is best considered a transcriptional regulatory hypomorph, even though its DNA-binding function is intact. The loss of oncogenic potential of EF DBD appears to be due to an underlying defect in transcriptional regulatory capability. This is an unanticipated result as the transcriptional regulation function of EWS/FLI was considered to be mediated solely by the EWS-portion of the fusion with the FLI-portion contributing only DNA-binding function.

### Capacity of EWS/FLI to mediate chromatin state is unaltered by deletions surrounding the FLI DNA-binding domain

It was recently reported that EWS/FLI functions as a pioneer transcription factor to open regions of chromatin that were previously closed (9, 15). As chromatin accessibility is a general necessity for transcriptional regulation, we next evaluated the role of EWS/FLI and its mutants on creation (or maintenance) of open chromatin states by performing ATAC-sequencing in our knock-down/rescue system. To focus on the role of the EWS/FLI mutants on chromatin accessibility, we overlapped the EWS/FLI-bound regions (as identified in our CUT&RUN analysis) with the ATAC-sequencing data. We found that ~95% of the nearly 13,000 EWS/FLI-binding sites had detectable ATAC signal (Figure 5A), indicating that most EWS/FLI binding peaks are associated with open chromatin states.

**Figure 5.**
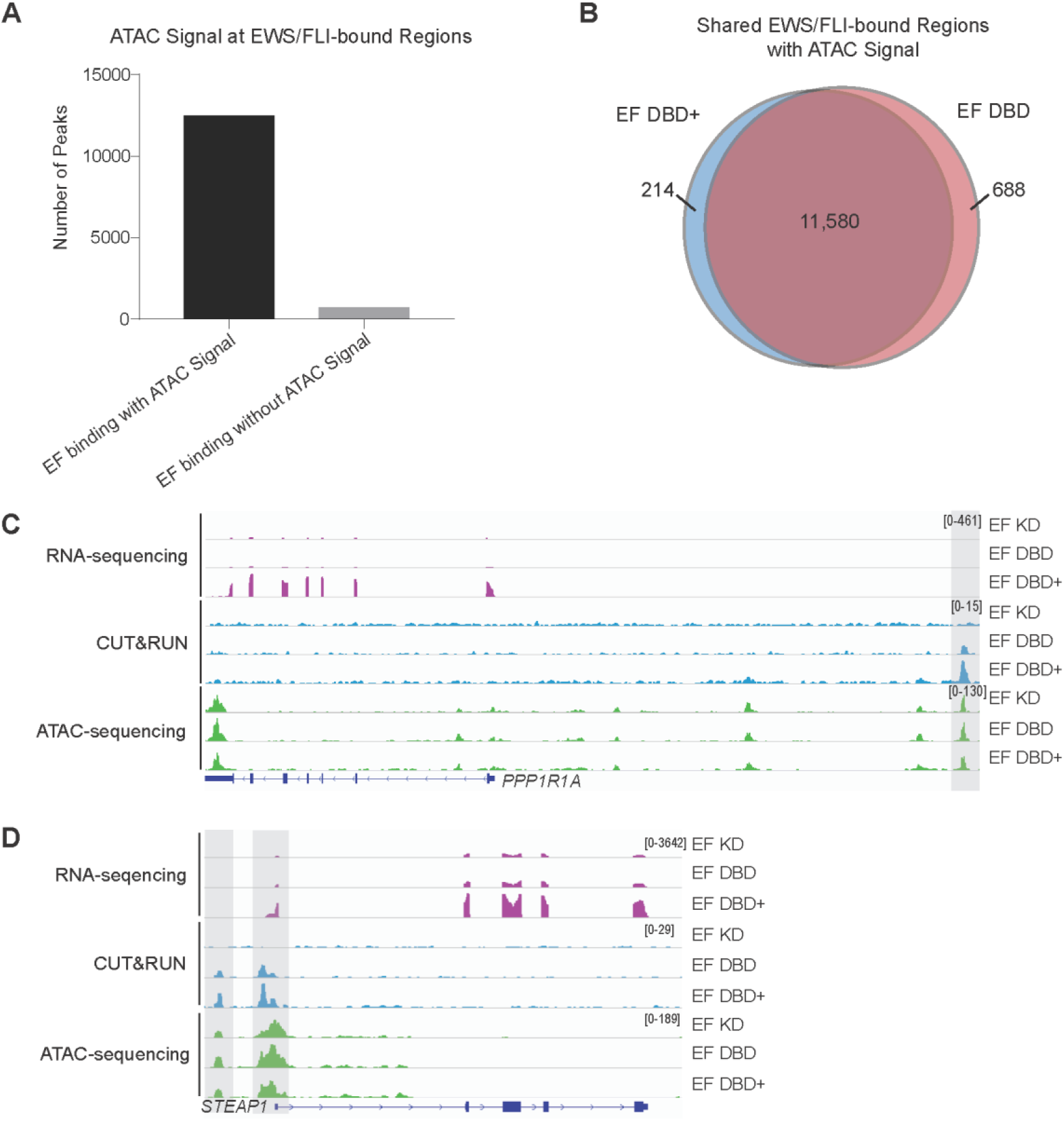
Chromatin-opening ability of EWS/FLI is unaltered by deletions flanking the FLI DNA-binding domain. (A) All EWS/FLI-bound loci in A673 cells (determined by CUT&RUN of knock-down/rescue cells expressing full-length EWS/FLI [EF]) were compared to loci harboring ATAC signal peaks and shown in graphical format. There were 12,482 EF-bound peaks with ATAC signal and 696 EF-bound peaks without ATAC signal. (B) Venn diagram analysis of regions bound by EF DBD+ and/or EF DBD that also had overlapping ATAC signals. (C-D) Representative tracks of RNA-sequencing, CUT&RUN genomic localization, and ATAC-sequencing signals for the indicated knock-down/rescue A673 cells (EF KD = iEF + EF; EF DBD = iEF + EF DBD; EF DBD+ = iEF + EF DBD+). Scales to view tracks were kept consistent across sequencing type in each panel and are represented on the right. Representative genes *PPP1R1A* (C) and *STEAP1* (D) are regulated by EF DBD+ but not EF DBD (adjusted p-value <0.05) and overlapping CUT&RUN and ATAC-sequencing peaks are highlighted.

To determine if EF DBD is defective in opening chromatin, we compared the ATAC signal at regions bound by EF DBD and those bound by EF DBD+. We found that almost 95% of ATAC peaks were shared between EF DBD+ and EF DBD (Figure 5B), suggesting that there were not significant differences in accessible chromatin associated with the two mutants.

To determine if more subtle differences in open chromatin might be associated with the capability of each mutant to regulate gene expression, we performed a heatmap analysis (Supplemental Figure 4A-B). At EWS/FLI-bound loci near genes that were regulated by EF DBD+ (but not EF DBD), we were surprised to find that the level of ATAC signal was similar in cells expressing EF DBD+ as compared to those expressing EF DBD. Interestingly, we also noted that the ATAC signal was similar at these sites in EWS/FLI knock-down cells (EF KD), indicating that the loss of EWS/FLI is not associated with a closing of the open chromatin state, at least in this system. Tracks comparing RNA-sequencing, CUT&RUN, and ATAC signal at representative genes only are shown in Figure 5C-D. These data indicate that the dysfunction of EF DBD in mediating gene regulation is not a consequence of an altered pioneer-type function to induce or maintain an open chromatin state at regulated genes.

### A fourth alpha-helix of the FLI ETS DNA-binding domain is essential for EWS/FLI-mediated oncogenic transformation

Finally, we sought to determine which flanking region of EF DBD+ (that is missing in EF DBD) is critical for oncogenic transformation. We engineered EF DBD+ constructs harboring deletions of either the amino-terminal 7-amino acids or the carboxyl-terminal 10-amino acids surrounding the core 85-amino acid DNA-binding domain of FLI (EF DBD+ ΔN or EF DBD+ ΔC, respectively; Figure 6A). Similar expression of each protein construct was observed in A673 cells utilizing our knock-down/rescue system (Figure 6B). Soft agar colony-forming assays demonstrated that EF DBD+ ΔN was fully-functional while EF DBD+ ΔC completely lost the ability to transform A673 cells (Figure 6C-D). These results clearly demonstrate that the 10-amino acids downstream of the FLI DNA-binding domain are essential for EWS/FLI-mediated oncogenesis.

**Figure 6.**
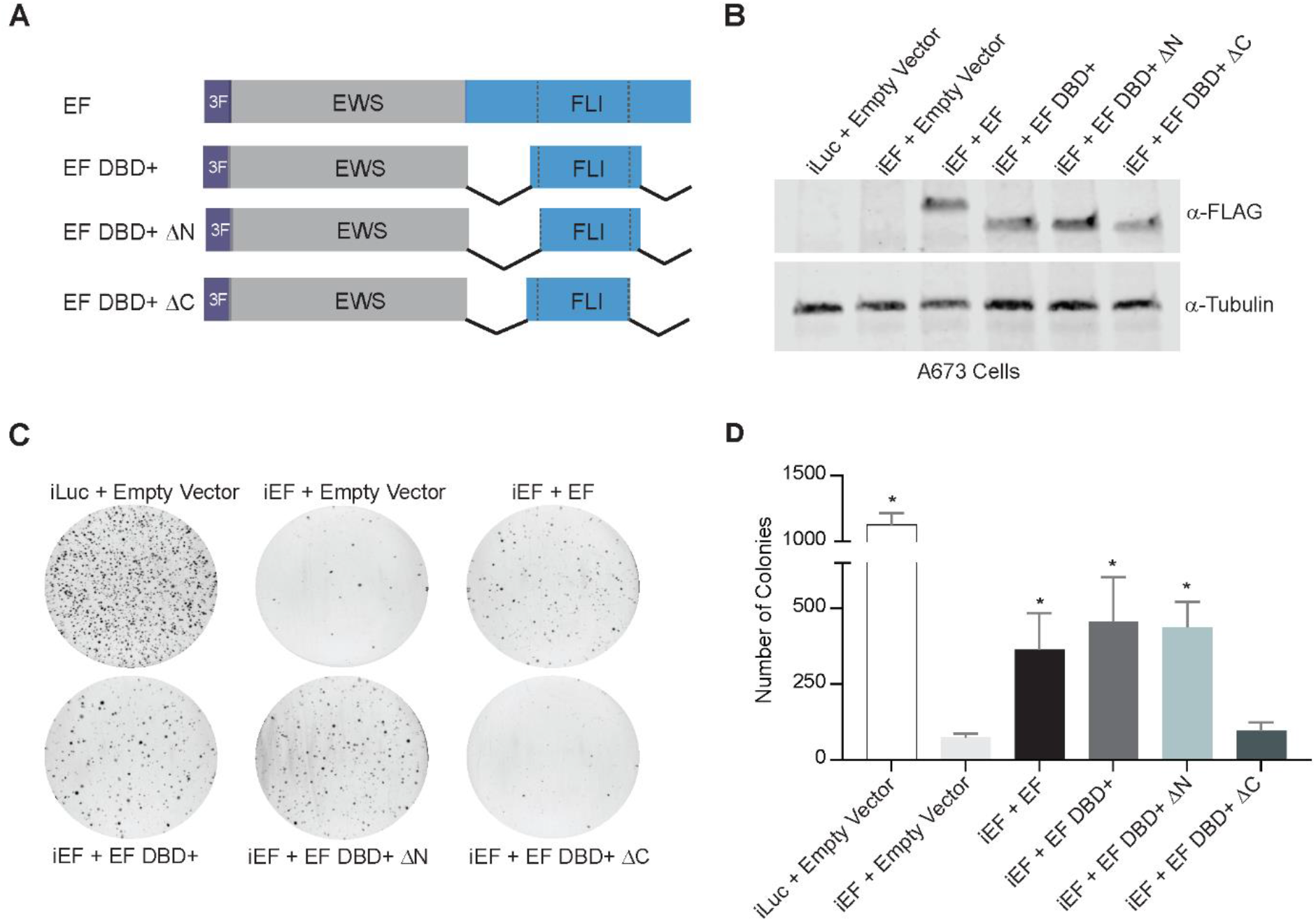
The carboxyl-terminal amino acids flanking the FLI DNA-binding domain are essential for EWS/FLI-mediated oncogenic transformation. (A) Protein schematics of 3xFLAG-tagged (3F) EWS/FLI constructs: EF represents a full-length EWS/FLI; EF DBD+ ΔN represents EWS/FLI containing the DBD+ version of FLI missing the 7-amino-terminal amino acids to the DBD; EF DBD+ ΔC represents EWS/FLI containing the DBD+ version of FLI missing the 10 carboxyl-terminal amino acids to the DBD. (B) Western blots of constructs expressed in A673 cells using our knock-down/rescue system. (C) Representative soft agar assay results are depicted for each of the indicated constructs. (D) Soft agar assay colony formation quantification. Data represented by mean ± SEM (n=3 independent replicates). Statistical significance of samples when compared to iEF + Empty Vector was used to determine the ability of each version of EWS/FLI to mediate oncogenic transformation in A673 cells. Asterisks indicate p-value < 0.05 as compared to negative control iEF + Empty Vector sample with no EWS/FLI expression.

Analysis of a previously published FLI protein crystal structure revealed that this 10-amino acid sequence forms two additional beta-turns and a fourth alpha-helix downstream of the winged helix-turn-helix DNA-binding domain of FLI (44). Interestingly, this same study demonstrated that recombinant FLI protein is capable of homodimerization when bound to an ETS HA site via interactions between this fourth alpha-helix of one FLI molecule with the a1-helix of another FLI molecule and that homodimerization was lost when a critical phenylalanine was mutated to alanine in this region (F362A) (44). We found that introduction of the F362A mutation to our EF DBD+ construct had no effect on oncogenic transformation in our A673 knock-down/rescue model system, indicating that homodimerization is not required for the oncogenic potential of EWS/FLI (Supplemental Figure 5A-D).

## Discussion

Ewing sarcoma represents a unique opportunity to understand how a single fusion protein mediates oncogenic transformation (8). The EWS/FLI fusion functions as an aberrant transcription factor to dysregulate the expression of several thousand genes and drive oncogenesis, but the mechanistic details of how EWS/FLI modulates this process are only beginning to be understood. It has long been recognized that the EWS-portion functions as a strong transcriptional activation domain (11). In recent years, it has become appreciated that this domain has unique biophysical characteristics that are important for its functions of self-association, recruitment of epigenetic regulators, and interaction with the basal transcriptional machinery that all cooperate to regulate gene expression (9, 14–17).

Although several studies suggested that the regions outside of the ETS DNA-binding domain of FLI may be important for EWS/FLI function, the FLI-portion of the fusion has largely been viewed as simply contributing DNA-binding function. In the current study, we took a systematic approach to understand the contributions of FLI to EWS/FLI activity in the context of a Ewing sarcoma cellular background. This allowed us to define a previously unappreciated role for the fourth alpha-helix of the extended FLI DNA-binding domain in transcriptional regulation. This alpha-helix does not appear to be important for the DNA-binding, genomic localization, or chromatin accessibility functions of EWS/FLI. Instead, loss of this helix results in a significant loss of gene-regulatory function that culminates in a complete loss of oncogenic transformation mediated by EWS/FLI.

The exact mechanism by which the fourth alpha-helix participates in gene regulation will require additional studies. One possibility is that the fourth alpha-helix is involved in protein-protein interactions with adjacent transcription factors. For example, it has been reported that EWS/FLI interacts with Serum Response Factor (SRF) on serum response elements to form a ternary complex with DNA that is required to regulate transcription (48). It was also shown that the AP-1 members, Fos-Jun, bind to AP-1 sites adjacent to ETS high-affinity sites and form a ternary complex with EWS/FLI (49). Although each of these interactions specifically occur with the FLI portion of EWS/FLI, they each occur outside of the regions of FLI contained in our EF DBD or EF DBD+ mutants and as such, do not readily explain the differences in activity observed between the proteins (49). EWS/FLI may interact with other transcription factors as well. However, we do not favor a loss of such EWS/FLI-transcription factor interactions as the most likely cause of the massive loss of transcriptional function by EF DBD. We reason that if there were losses of EWS/FLI interactions with specific transcription factors, we would have expected a more limited loss of gene expression (rather than the ~70% loss we observed for EF DBD). Furthermore, the formation of ternary complexes between pairs of transcription factors with DNA tend to stabilize DNA binding. As such, we might also have anticipated a significant change in genomic localization of EF DBD, which was not observed. We currently favor a model whereby the fourth alpha-helix interacts with epigenetic regulators, and/or components of the core transcriptional machinery, that are required for global gene regulation, rather than regulation limited to specific loci.

Work in NIH3T3 mouse fibroblasts suggested a role for the carboxyl-terminal region of FLI in mediating transcriptional down-regulation by EWS/FLI (31). Our work here rules out a significant role for this region in EWS/FLI-mediated oncogenesis. Additionally, luciferase reporter assays have long been used as functional screens, but our results show that activation on a luciferase reporter in an artificial system does not necessarily translate to a Ewing sarcoma model. Indeed, we also note that we did not see direct evidence of the pioneer-type function of EWS/FLI in the Ewing sarcoma model, which had primarily been observed in a mesenchymal stem cell model (9). In our system, EWS/FLI-occupied sites remained open and accessible following knock-down of EWS/FLI. It may be that the 80-90% knock-down we achieved was insufficient to allow for chromatin closing of those loci, or perhaps insufficient time was provided to allow for chromatin closing. Nevertheless, changes in chromatin accessibility were not associated with the transcriptional dysfunction exhibited by EF DBD. These findings highlight the importance of analyzing EWS/FLI activity in a relevant Ewing sarcoma cellular context.

It is important to note that a detailed comparison of protein structures of ETS family members revealed that many of these proteins harbor this additional fourth alpha-helix just downstream of their DNA-binding domains. Therefore, the work presented here may have relevance beyond the EWS/FLI-associated Ewing sarcoma context. For example, Ewing sarcoma translocations involve one of five closely-homologous ETS family members (FLI, ERG, FEV, ETV1, and ETV4) (11). Additionally, *TMPRSS2-ERG* fusions exist in approximately 50% of prostate cancer cases, with *TMPRSS2-FEV, -ETV1, -ETV4*, and *-ETV5* fusions found in other patients (50). In fact, many ETS family members have been implicated in the oncogenesis of numerous solid and liquid tumor types via mechanisms of over-expression, amplification, mutations, and translocations (20). As the functional motif we identified as crucial for EWS/FLI activity is conserved in numerous other ETS factors, the data presented in this report may have wide-ranging implications for oncogenesis in multiple tumor types.

## Conclusions

In summary, we have taken a systematic structure-function approach to identify a previously unappreciated region in the extended FLI DNA-binding domain that is required for transcriptional regulation and oncogenic transformation mediated by EWS/FLI. This transcriptional function is distinct from the DNA-binding and genomic localization functions typically associated with the ETS domain. This work has implications not only for the development of Ewing sarcoma, but may also be useful in understanding the development of other ETS-associated tumors and, perhaps, even normal ETS transcriptional function. A better understanding of this newly-defined region may lead to novel approaches for therapeutically-targeting EWS/FLI, as well as other ETS factors. Ultimately, these efforts may lead to more efficacious therapeutic options for patients with this devastating disease.

## Abbreviations

ATAC-sequencing: Assay for Transposase-Accessible Chromatin using sequencing
CUT&RUN: Cleave Under Targets & Release Under Nuclease
cDNA: complementary DNA
DBD: DNA-binding domain
DNA: deoxyribonucleic acid
EF: experimental EWS/FLI cDNA constructs
ERG: ETS-related gene
ETS: E26 transformation specific
ETV1: ETS variant transcription factor 1
ETV4: ETS variant transcription factor 4
ETV5: ETS variant transcription factor 5
*EWSR1* (EWS): Ewing sarcoma breakpoint region 1
FET: FUS/TLS, EWSR1, TAF15
FEV: Fifth Ewing variant protein
*FLI1* (FLI): Friend leukemia integration 1
FUS: Fused in sarcoma
GSEA: Gene Set Enrichment Analysis
HA: high-affinity
HEK-293EBNA: Human embryonic kidney-293 cell line expressing Epstein Barr nuclear antigen
IGV: Integrated Genome Viewer
KD: knock-down
K_D_: dissociation constant
log2(FC): log2(Fold Change)
mRNA: messenger RNA
μSat: microsatellite
NES: Normalized Enrichment Score
qRT-PCR: quantitative Reverse Transcriptase-Polymerase Chain Reaction
RNA: ribonucleic acid
SEM: standard error of the mean
SRF: Serum Response Factor
STR: short tandem repeats
TMPRSS2: Transmembrane protease, serine 2

## Declarations

### Ethics Approval and Consent to Participate

Not applicable.

## Consent for publication

Not applicable.

## Availability of data and materials

The sequencing datasets generated and analyzed during the current study are available in the Gene Expression Omnibus and accessible at [unique identifier number pending]. All other data generated or analyzed during this study are available from the corresponding author on reasonable request.

## Competing interests

SLL declares a competing interest as a member of the advisory board for Salarius Pharmaceuticals. SLL is also a listed inventor on United States Patent No. US 7,939,253 B2, “Methods and compositions for the diagnosis and treatment of Ewing’s sarcoma,” and United States Patent No. US 8,557,532, “Diagnosis and treatment of drug-resistant Ewing’s sarcoma.” This does not alter our adherence to *Molecular Cancer* policies on sharing data and materials.

## Funding

Research reported in this publication was supported by the National Institute of Health award T32 GM068412 to MAB, and U54 CA231641 to SLL. The content is solely the responsibility of the authors and does not necessarily represent the official views of the National Institutes of Health.

## Authors Contributions

MAB and SLL are responsible for conceptualization of the project. Investigation was performed by MAB, JCC, JSA, BDS, and BZS. Methodology was formulated by MAB, JSA, ERT, IS, and BZS. Data analysis was performed by MAB, CT, ERT, IS, and MW. Manuscript preparation was completed by MAB and reviewing and editing was performed by all authors. Funding acquisition was completed by MAB and SLL. Supervision was provided by SLL.

## Acknowledgements

We thank Dr. Andrea K. Byrum, Dr. Kirsten N. Johnson, Dr. Kathleen I. Pishas, Dr. Jack Tokarsky, and Ariunaa Bayanjargal for thoughtful discussion concerning the hypothesis and methodology of this manuscript.

---

**Supplemental Figure 1.**
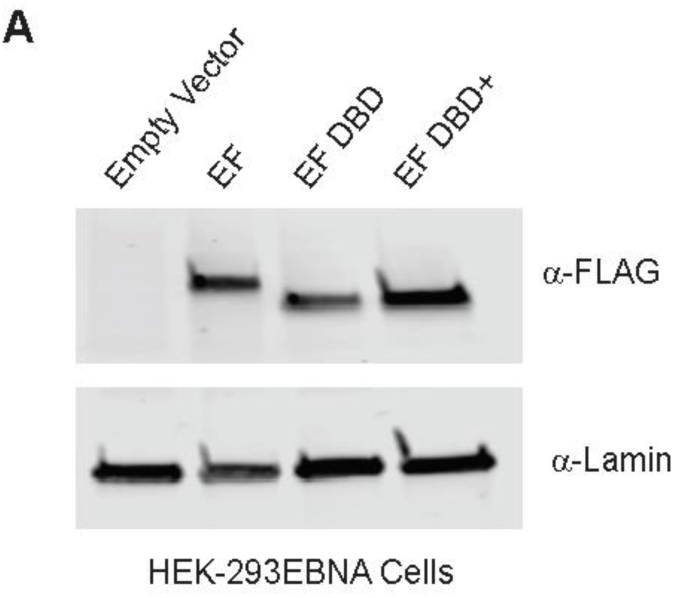
EWS/FLI mutant construct expression in HEK-293EBNA cells. (A) 3xFLAG-tagged full-length EWS/FLI (EF), EF DBD, or EF DBD+ constructs were expressed in HEK-293EBNA cells. Western blot analysis was used to determine expression of these proteins utilizing α-FLAG. α-Lamin was used as a loading control.

**Supplemental Figure 2.**
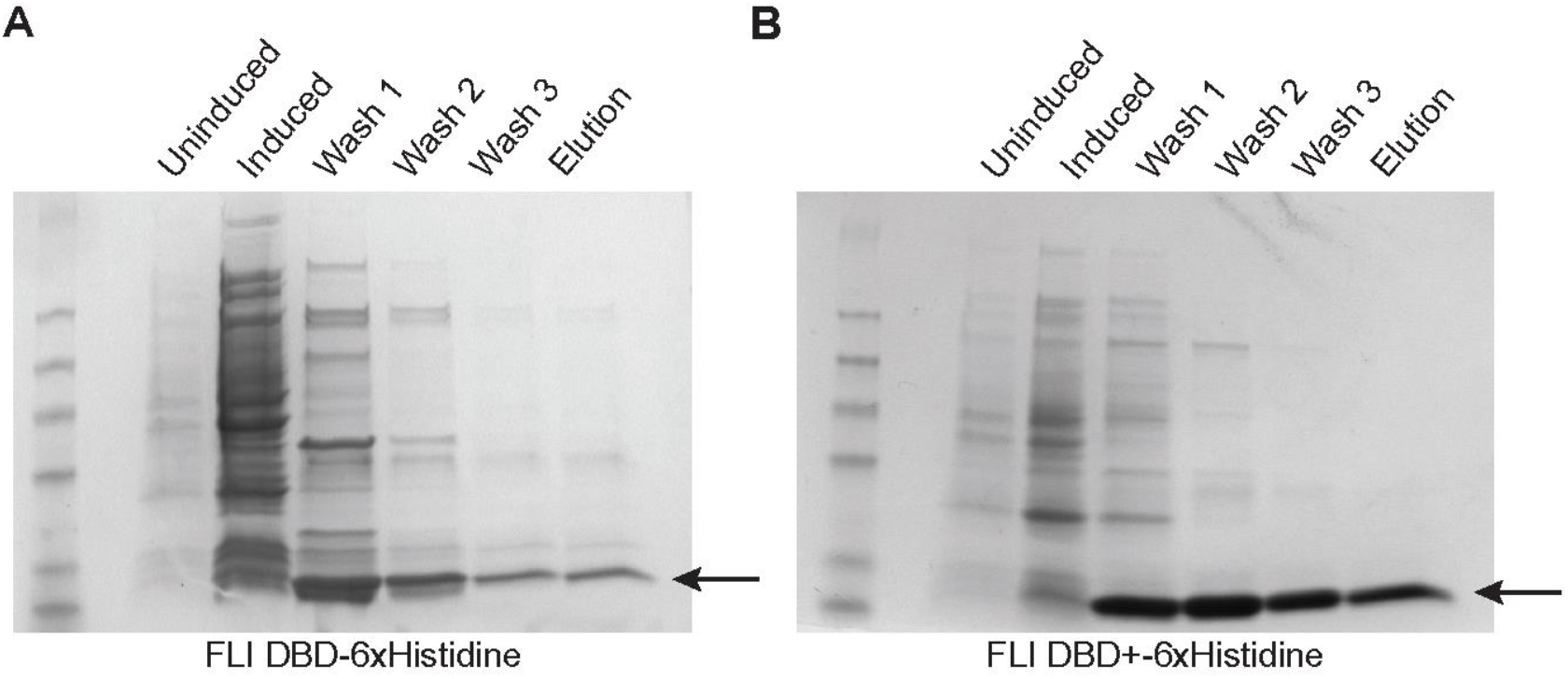
Recombinant FLI DBD and FLI DBD+ protein purification. (A-B) Samples were taken at several stages of recombinant protein purification for FLI DBD-6xHisitidine (A) and FLI DBD+-6xHistidine (B), including: Uninduced bacteria, Induced bacteria, after Wash 1, Wash 2, and Wash 3 following bacterial lysis, and Eluted Fraction. Samples were run on a SDS-PAGE gel and demonstrate good purity at the final elution step.

**Supplemental Figure 3.**
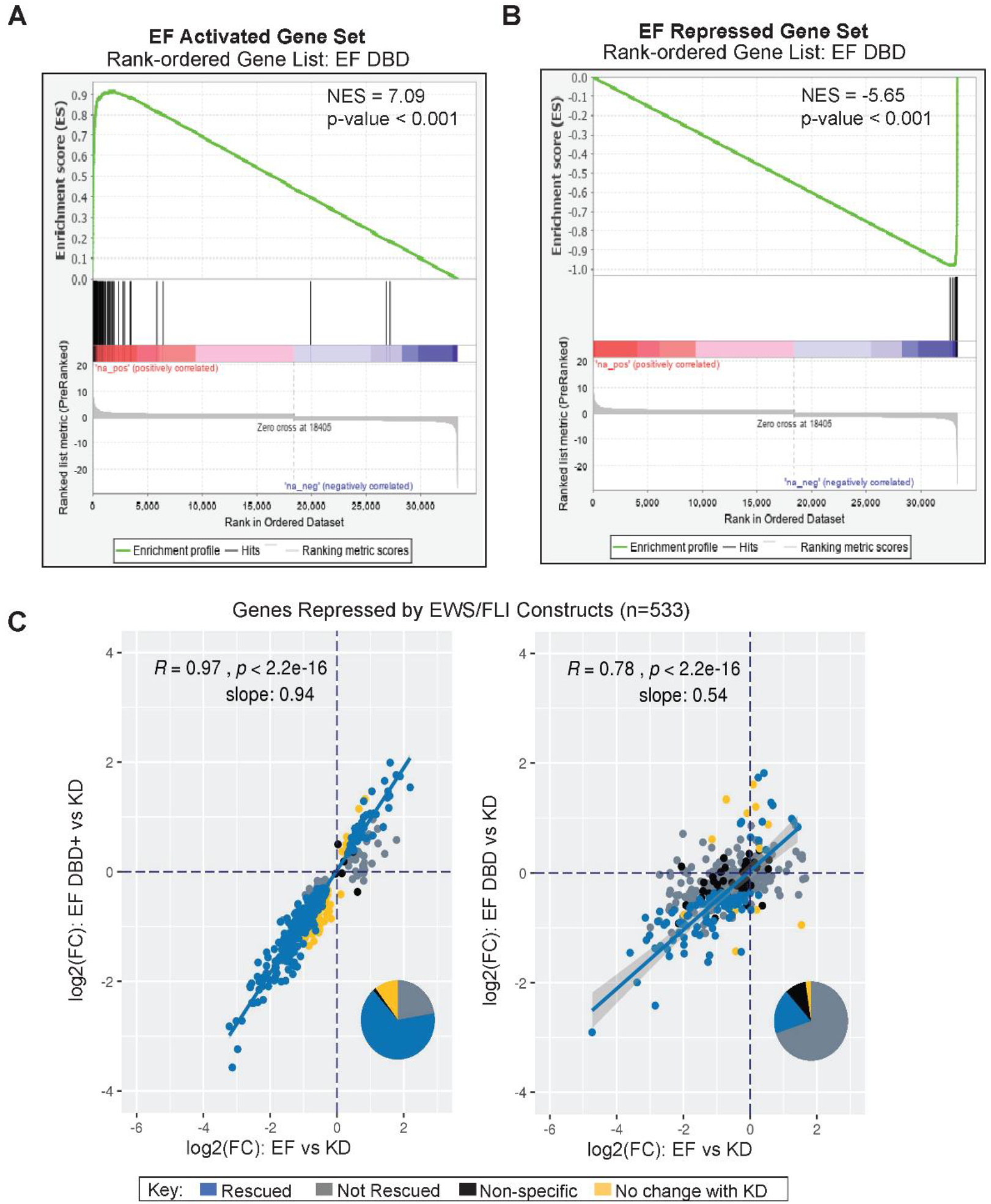
Deletions surrounding the FLI DBD of EWS/FLI result in weaker transcriptional regulation. (A-B) GSEA analysis comparing genes regulated by EF DBD as the rank-ordered gene list to (A) a gene set activated by EF (log2(FC) > 1.5, FDR < 0.05) or (B) a gene set repressed by EF (log2(FC) < −1.5, FDR < 0.05). (C) Genes significantly repressed by endogenous EWS/FLI were defined using a previous RNA-sequencing dataset (47). Genes repressed by EF, EF DBD, and EF DBD+ were compared to this list of EWS/FLI-repressed genes. Scatterplots comparing genes repressed by exogenous EF (on the x-axis) to EF DBD+ (left) or EF DBD (right) (on the y-axis) were plotted to determine the ability of these constructs to rescue repression of these genes. Significance was defined by a log2(FC) < 0 and an adjusted p-value < 0.05. Pearson correlation coefficient and associated p-values with slope are noted on the plots. Pie charts represent the proportion of genes found in each of the described groups is also depicted.

**Supplemental Figure 4.**
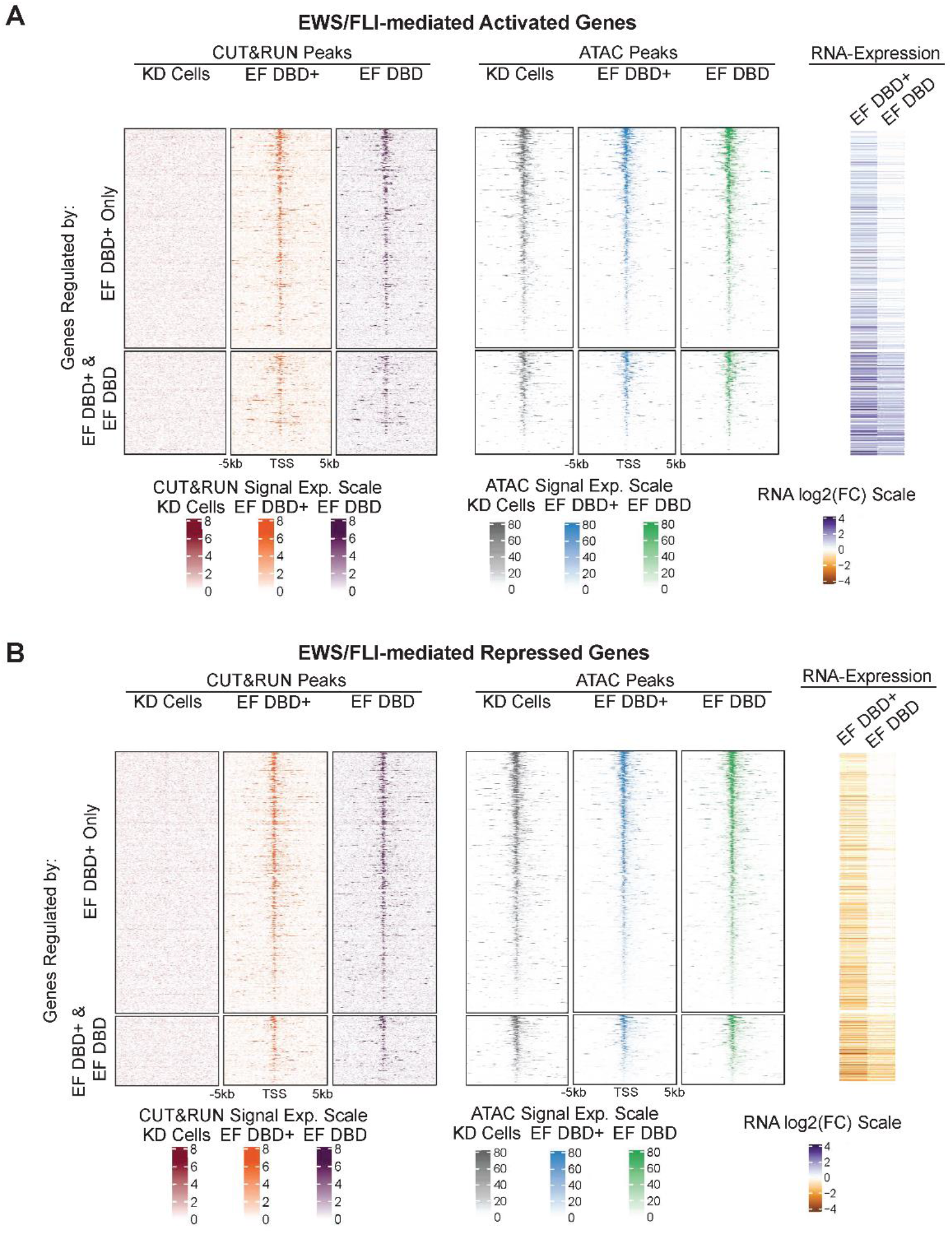
Heatmap analysis of ATAC-signal at EWS/FLI-mutant bound activated and repressed genes. (A-B) Heatmaps depicting CUT&RUN and ATAC-sequencing signals, centered on the nearest transcriptional start sites (TSS) of genes regulated by EF DBD+ only or both EF DBD+ and EF DBD. EWS/FLI-mediated activated genes are visualized in (A) and repressed genes in (B). Knock-down cells (KD; iEF + Empty Vector), EF DBD+ (iEF + EF DBD+), and EF DBD (iEF + EF DBD) were compared (scales for peak height are depicted below heatmaps). The log2(FC) of RNA expression for EF DBD+ and EF DBD (compared to KD) is pictured on the right with log2(FC) scale depicted below.

**Supplemental Figure 5.**
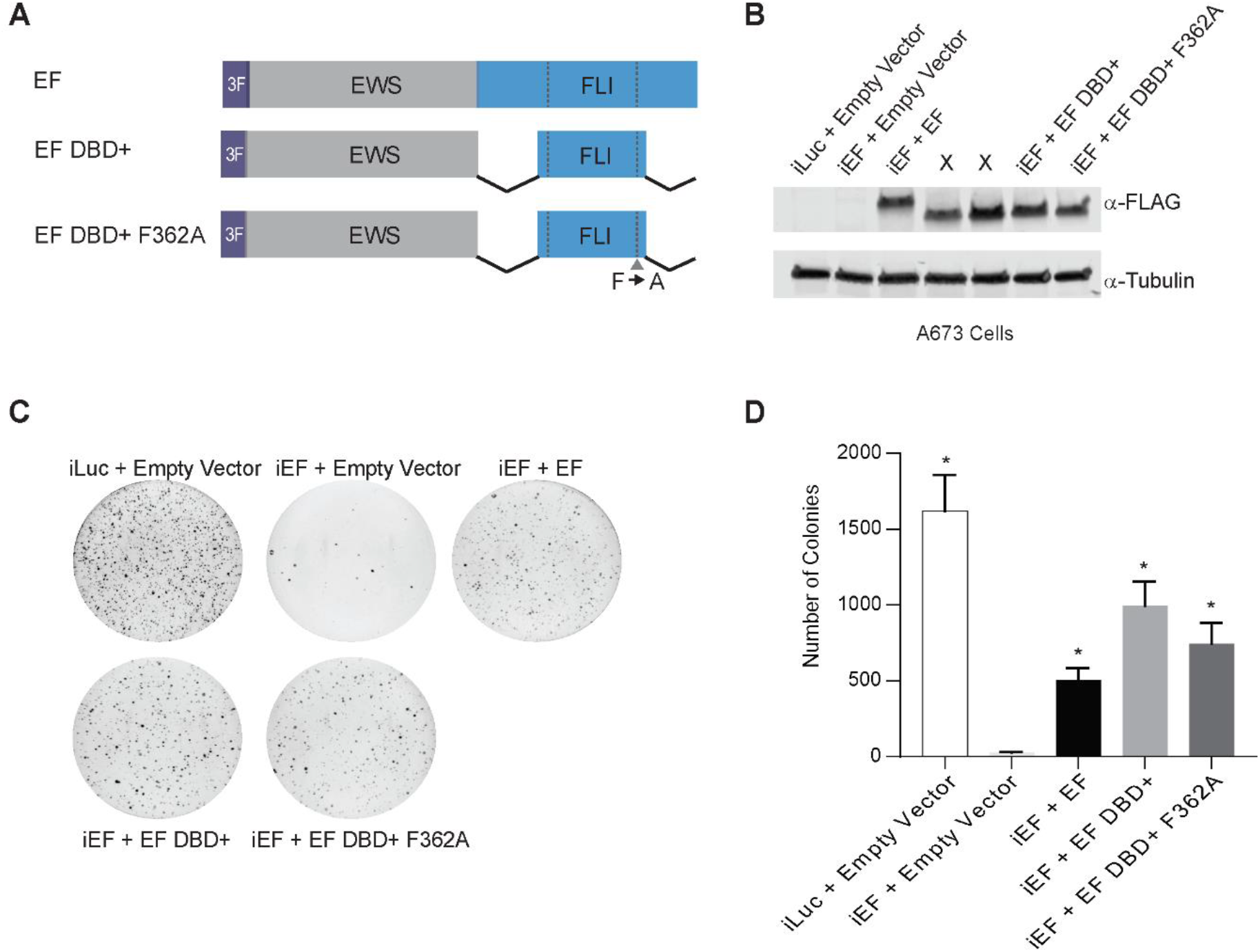
Homodimerization motif is dispensable for EWS/FLI-mediated oncogenic transformation. (A) Protein schematics of 3xFLAG-tagged (3F) EWS/FLI constructs: EF represents full-length EWS/FLI; EF DBD+ contains EWS fused to the DBD+ version of FLI; EF DBD+ F362A represents EWS fused to the DBD+ version of FLI with the phenylalanine “F” residue at residue 362 mutated to alanine “A”. (B) Constructs were expressed in A673 cells using our knock-down/rescue system and Western blot analysis was used to determine efficient expression of these proteins using α-FLAG antibody for detection of EWS/FLI constructs and α-Tubulin as a loading control. (Samples labeled as X are not relevant to this current set of experiments.) (C) Representative soft agar assay results are depicted, including controls and experimental samples. (D) Soft agar assay colony formation quantification. Data is represented by mean ± SEM (n=3 independent replicates). Asterisks indicate p-value <0.005 as compared to samples with EWS/FLI knock-down (iEF + Empty Vector).

**Additional file 1.** Amino acids references for EWS/FLI Mutant Constructs

| **Construct Name** | **Amino acids residues according to protein:** |  |  |
| --- | --- | --- | --- |
|  | **EWSR1**<br>(NP_001156757.1) | **FLI1**<br>(NP_002008.2) | **“Type IV” EWS/FLI**<br>(May *et al.*, 1993) |
| Full-length “Type IV” EF | 1-265 | 242-452 | 266-476 |
| EF ΔN-FLI | 1-265 | 275-452 | 299-476 |
| EF ΔC-FLI | 1-265 | 242-373 | 266-397 |
| EF DBD | 1-265 | 277-361 | 302-386 |
| EF DBD+ | 1-265 | 270-371 | 295-396 |
| FLI DBD | - | 277-361 | 302-386 |
| FLI DBD+ | - | 270-371 | 295-396 |
| EF DBD+ ΔN | 1-265 | 277-371 | 302-396 |
| EF DBD+ ΔC | 1-265 | 270-361 | 295-386 |
| EF DBD+ F362A | 1-265 | 270-371, F362A | 295-396, F362A |

**Additional file 2.**
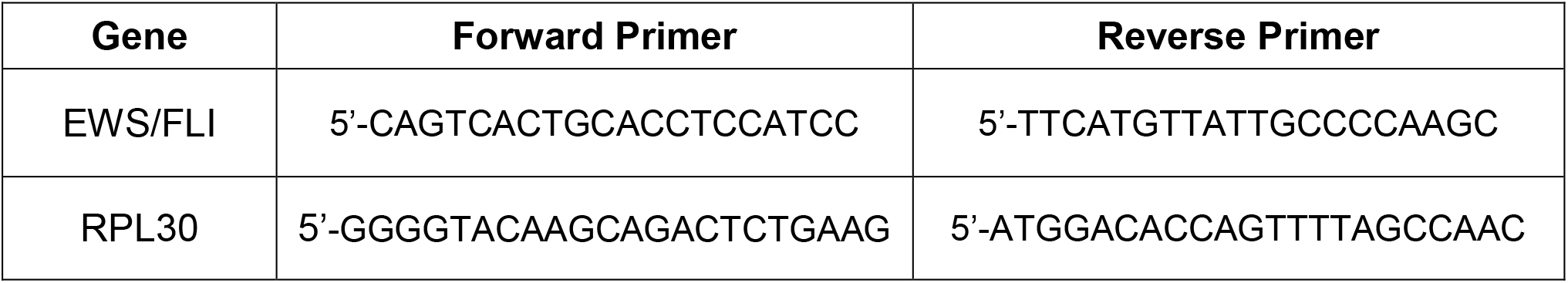
Sequences for primers used in RT-qPCR experiments

**Additional file 3.**
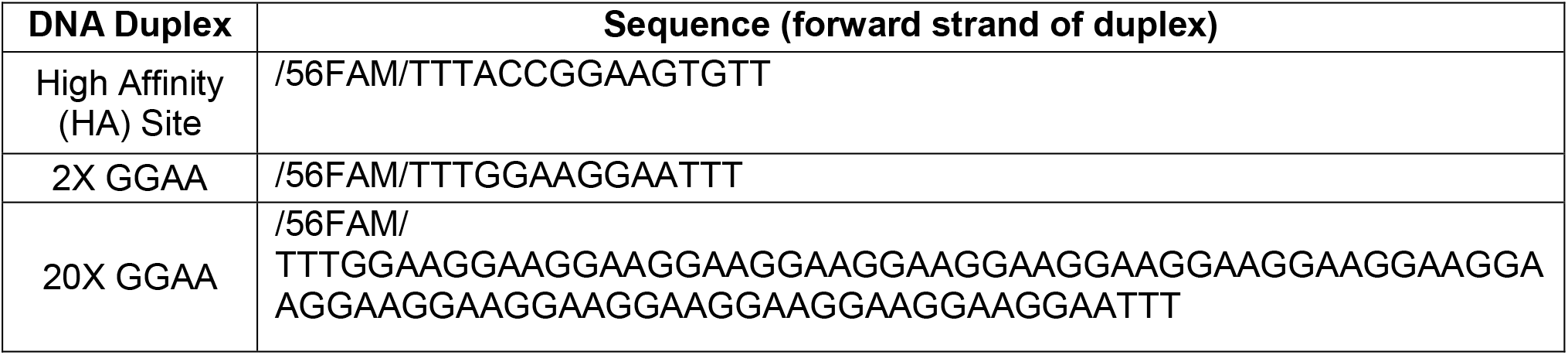
Fluorescein-labelled DNA-duplex oligonucleotides used for fluorescence anisotropy experiments

